# *Peromyscus leucopus*, *Mus musculus*, and humans have distinct transcriptomic responses to larval *Ixodes scapularis* bites

**DOI:** 10.1101/2024.05.02.592193

**Authors:** Jeffrey S. Bourgeois, Julie E. McCarthy, Siu-Ping Turk, Quentin Bernard, Luke H. Clendenen, Gary P. Wormser, Luis A. Marcos, Kenneth Dardick, Sam R. Telford, Adriana R. Marques, Linden T. Hu

## Abstract

*Ixodes scapularis* ticks are an important vector for at least six tick-borne human pathogens, including the predominant North American Lyme disease spirochete *Borrelia burgdorferi*. The ability for these ticks to survive in nature is credited, in part, to their ability to feed on a variety of hosts without excessive activation of the proinflammatory branch of the vertebrate immune system. While the ability for nymphal ticks to feed on a variety of hosts has been well-documented, the host-parasite interactions between larval *I. scapularis* and different vertebrate hosts is relatively unexplored. Here we report on the changes in the vertebrate transcriptome present at the larval tick bite site using the natural *I. scapularis* host *Peromyscus leucopus* deermouse, a non-natural rodent host *Mus musculus* (BALB/c), and humans. We note substantially less evidence of activation of canonical proinflammatory pathways in *P. leucopus* compared to BALB/c mice and pronounced evidence of inflammation in humans. Pathway enrichment analyses revealed a particularly strong signature of interferon gamma, tumor necrosis factor, and interleukin 1 signaling at the BALB/c and human tick bite site. We also note that bite sites on BALB/c mice and humans, but not deermice, show activation of wound-healing pathways. These data provide molecular evidence of the coevolution between larval *I. scapularis* and *P. leucopus* as well as expand our overall understanding of *I. scapularis* feeding.

**Significance:** *Ixodes scapularis* tick bites expose humans to numerous diseases in North America. While larval tick feeding enables pathogens to enter the tick population and eventually spread to humans, how larval ticks interact with mammals has been understudied compared to other tick stages. Here we examined the transcriptomic response of a natural *I. scapularis* rodent host (*Peromyscus leucopus*), a non-native *I. scapularis* rodent host (*Mus musculus*), and an incidental host (humans). We find that there are differences in how all three species respond to larval *I. scapularis*, with the natural host producing the smallest transcriptomic signature of a canonical proinflammatory immune response and the incidental human host producing the most robust signature of inflammation in response to the larval tick. These data expand our understanding of the pressures on ticks in the wild and inform our ability to model these interactions in laboratory settings.

## Introduction

*Ixodes scapularis* (formerly *Ixodes dammini*) ticks are the most important invertebrate vector of human diseases in North America [1]. These ticks are responsible for spreading most cases of Lyme disease (predominantly caused by *Borrelia burgdorferi* sensu stricto in North America), which affects roughly 476,000 individuals in the US yearly [2], as well as six other human pathogens—*Anaplasma phagocytophilum*, *Babesia microti*, *Borrelia miyamotoi*, *Borrelia mayonii*, *Ehrlichia muris eauclairensis* and deer tick virus/Powassan virus [3, 4]. The spread of pathogens into and out of *I. scapularis* requires a variety of complex interactions to occur at the skin-vector interface [1, 5]. Tick feeding can trigger a immunological processes which threaten the ability for the tick to survive the blood meal [6]. To prevent this, *Ixodes* ticks secrete saliva into the feeding site, exposing the host to anticoagulants and immunomodulatory compounds, which dampens the host immune response and ensures the tick remains attached until completion of feeding [7–9]. This secretion also contributes to the spread of *B. burgdorferi* into new hosts—both by providing a mechanism to exit the tick [10–12] and by dampening the immune response while the pathogen establishes the infection [13–20].

*I. scapularis* is a generalist parasite that successfully feeds on many hosts [21]. In line with this, previous work has demonstrated that nymphal tick bites on guinea pigs (*Cavia porcellus*), *Mus musculus*, *Peromyscus leucopus*, and humans are broadly similar at the histopathological level [22–24]. However, major changes across species occur during recurrent tick bites, where, for instance, guinea pigs and humans become substantially more inflamed than the other species—resulting in itching, rejection of the tick, and/or reduced risk of pathogen transmission [22, 24, 25]. Notably, at the transcriptional level, differences between *M. musculus* (BALB/c) and *C. porcellus* immune responses to nymphal *I. scapularis* were apparent even at the first feeding [24]. This demonstrates that not all *I. scapularis* hosts are equally permissive to parasitization.

Relatively little attention has been paid to how any vertebrate—including humans or rodents—interact with larval *I. scapularis*, though field studies have documented a strong association between larvae and *P. leucopus* in the northeastern and midwestern United States [21, 26–28], as well as birds and reptile hosts in the southeastern United States [21, 29, 30]. In his 1989 review, Ribeiro stated that their unpublished data demonstrated that larval *I. scapularis* can feed efficiently on the North American deermouse *P. leucopus* but not on *C. porcellus* [6]. Similarly, larval *I. scapularis* were found to feed better on *P. leucopus* than *Microtus pennsylvanicus* voles [31]. Larval *I. scapularis* are not infected with *B. burgdorferi* [32] and less commonly bite humans compared with the nymphal stage [33]; thus they have very limited direct clinical impact on patient health. However, this stage does have a major indirect impact on human health by impacting *B. burgdorferi* abundance in nature—larval tick feeding is critical for the continuation of the *B. burgdorferi* enzootic cycle [1].

In this study, we compare the transcriptomic response of two rodent models of tick feeding, the natural *I. scapularis* host *P. leucopus* and the artificial *I. scapularis* host *M. musculus* (BALB/c), to examine the transcriptomic response to larval *I. scapularis*. Pathway enrichment analyses revealed activation of more proinflammatory signaling pathways in *M. musculus* than in *P. leucopus*. We also note evidence of tissue remodeling and homeostasis genes activating in BALB/c rodents but not *P. leucopus*. Integration of a previously published transcriptomic dataset examining the nymphal bite site in BALB/c mice demonstrated considerable differences between the *M. musculus* immune response to each of these tick stages, suggesting that ticks may have different capacities to suppress proinflammatory pathways in different hosts across the stages in their life cycle. We also evaluated human patient samples [34] to compare how the human transcriptional response compared to these rodent responses. We found a strong proinflammatory immune response to larval *I. scapularis* in humans, with activation of many of the same pathways (tumor necrosis factor (TNF), interferon gamma (IFN-γ), interleukin 1 (IL1) signaling) as in *M. musculus*. Overall, this study substantially increases our understanding of host-*Ixodes* interactions.

## Results

### BALB/c mice and *Peromyscus leucopus* deermice respond differently to larval *Ixodes scapularis*

In order to examine how BALB/c (n=4) and *P. leucopus* (n=5) respond to larval *I. scapularis* bites, 10 larval ticks were placed in a tick containment chamber affixed to the back of each rodent. Because larval ticks do not leave a noticeable bite site after detachment on either rodent, 2mm punch biopsies were taken surrounding a feeding tick at 48 hours post-placement roughly one-half to three quarters of the way through feeding. Additionally, a second punch biopsy was taken from a region outside the tick containment chamber as a no tick bite control (**Figure 1A**).

**Figure 1:**
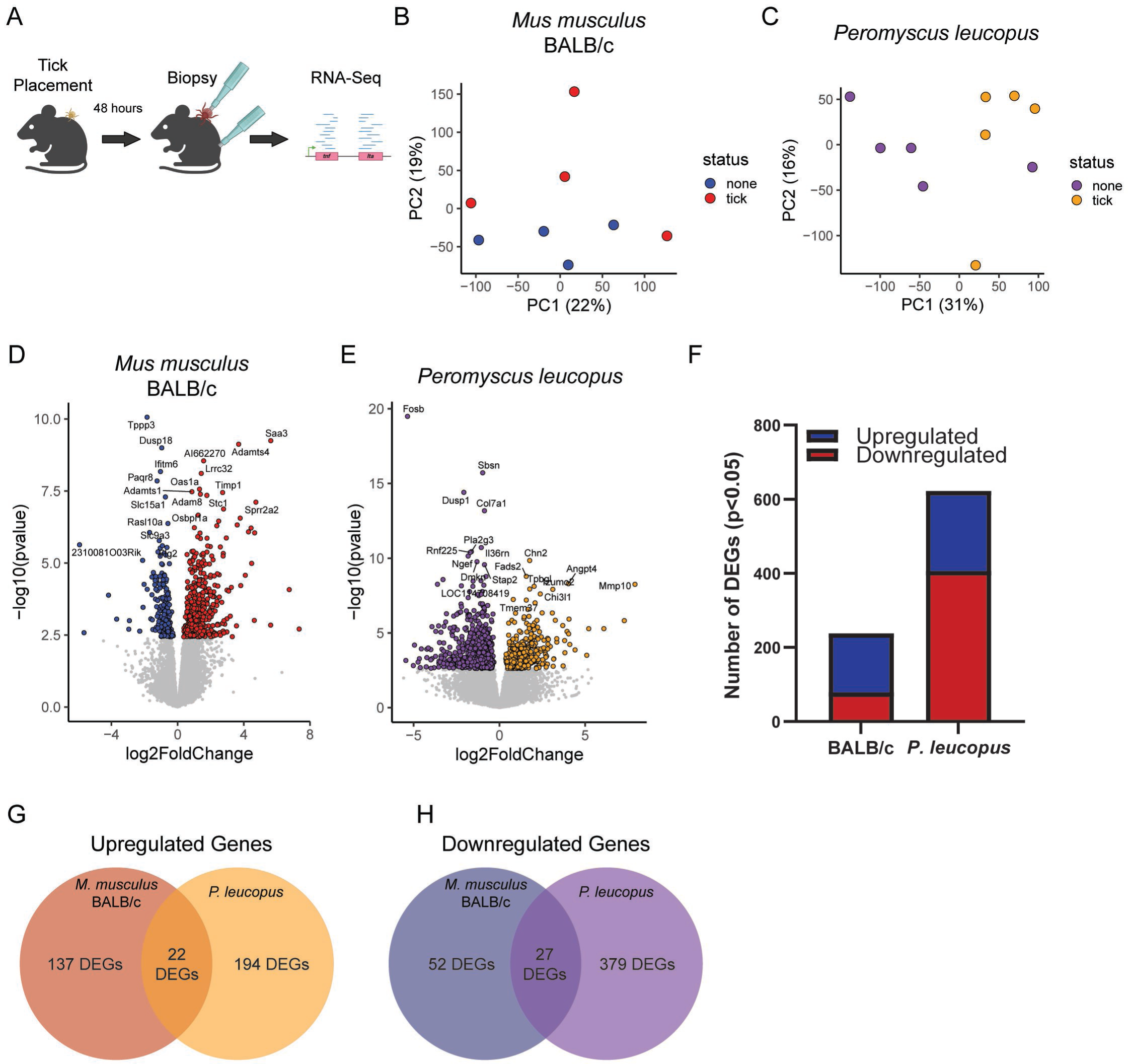
BALB/c *M. musculus* and *P. leucopus* display different transcriptomic responses to bites from larval *I. scapularis*. (A) Schematic of larval tick placement and RNA-seq. (B,C) Principal component analysis based on observed transcripts per million reads for each gene in (B) BALB/c mice or (C) *P. leucopus*. (D,E) Volcano plots of differentially expressed genes in (D) BALB/c mice or (E) *P. leucopus*. Genes with p>0.05 shown in gray. (F) BALB/c mice display more upregulation of genes than downregulation, while *P. leucopus* display more downregulation of genes than upregulation. (G, H) Most genes (G) upregulated or (H) downregulated in BALB/c or *P. leucopus* are not similarly differentially expressed in the opposite rodent.

We performed RNA sequencing on these biopsies. Principal component analysis revealed that tick bite sites cluster away from unbitten skin along the second principal component in BALB/c mice (**Figure 1B**) and the first principal component in *P. leucopus* (**Figure 1C**). We next performed differential gene expression analysis (**Table S1**) to examine genes that were induced or suppressed (p_adj_<0.05) by larval tick feeding. For all analyses, gene lists were restricted to those genes that could be reliably identified across all samples—meaning across different skin biopsy sites (tick bite and control) and across species (BALB/c and *P. leucopus*). Differential gene expression analyses revealed that 238 genes were differentially expressed in BALB/c tick bite sites (**Figure 1D**), and 622 genes were differentially expressed in *P. leucopus* tick bite sites (**Figure 1E**). Examining the direction of effect, we found that most differentially expressed genes in BALB/c were upregulated (159 upregulated, 79 downregulated), while most genes in *P. leucopus* were downregulated (216 upregulated, 406 downregulated) (**Figure 1F**). There was remarkably little overlap between the upregulated genes or downregulated genes across BALB/c mice and *P. leucopus* (**Figures 1G, 1H**).

### Ingenuity Pathway Analysis reveals more proinflammatory cytokine signaling in BALB/c mice compared to *P. leucopus* at tick bite sites

The Qiagen Ingenuity Pathway Analysis (IPA) pipeline [35] can be used to identify signaling pathways that show evidence of activation or suppression in response to the tick bite. This program assigns pathways controlled by a given upstream regulator (*e.g.* cytokines, transcription factors). Importantly, this is based on gene expression of genes across the entire pathways, and thus these values cannot be interpreted as meaning that the upstream regulator itself (*e.g. Tnf*) is upregulated or downregulated in each dataset.

Using this tool, we identified 48 cytokine-regulated pathways that show evidence (Z-score ≥ 2) of activation and one pathway that showed evidence (Z-score ≤ −2) of suppression in BALB/c (**Table S2**). In contrast three cytokine-regulated pathways showed evidence of activation and two showed evidence of suppression in *P. leucopus* (**Table S2**). Of these, two pathways (IFNB1 and PRL) were activated in both species. We focused our attention on the 23 cytokine-regulated pathways that had (1) the most significant predicted activation (Z-score > 3) or repression (Z-score < −3) in at least one of the species, and (2) had at least a Z-score difference of 2 between the two species (**Figure 2A**). This revealed multiple proinflammatory pathways that were predicted to have substantial activation in BALB/c mice but not *P. leucopus*, particularly the tumor necrosis factor (TNF) (BALB/c Z-score 6.132, *P. leucopus* Z-score −0.867) and interferon-gamma (IFNG) regulated pathways (BALB/c Z-score 6.4, *P. leucopus* Z-score - 0.02). We also note increased predicted IL6 and IL1 signaling in BALB/c mice but not *P. leucopus*. Conversely, BALB/c mice had substantial predicted repression of the anti-inflammatory interleukin 1 receptor antagonist (IL1RN) regulated pathway, which showed no evidence of regulation in *P. leucopus* (BALB/c Z-score −3.633, *P. leucopus* Z-score 0.454). Together, these data suggest that the BALB/c larval *I. scapularis* shows signs of robust proinflammatory cytokine signaling at the transcriptomic level that is substantially less present in *P. leucopus*.

**Figure 2:**
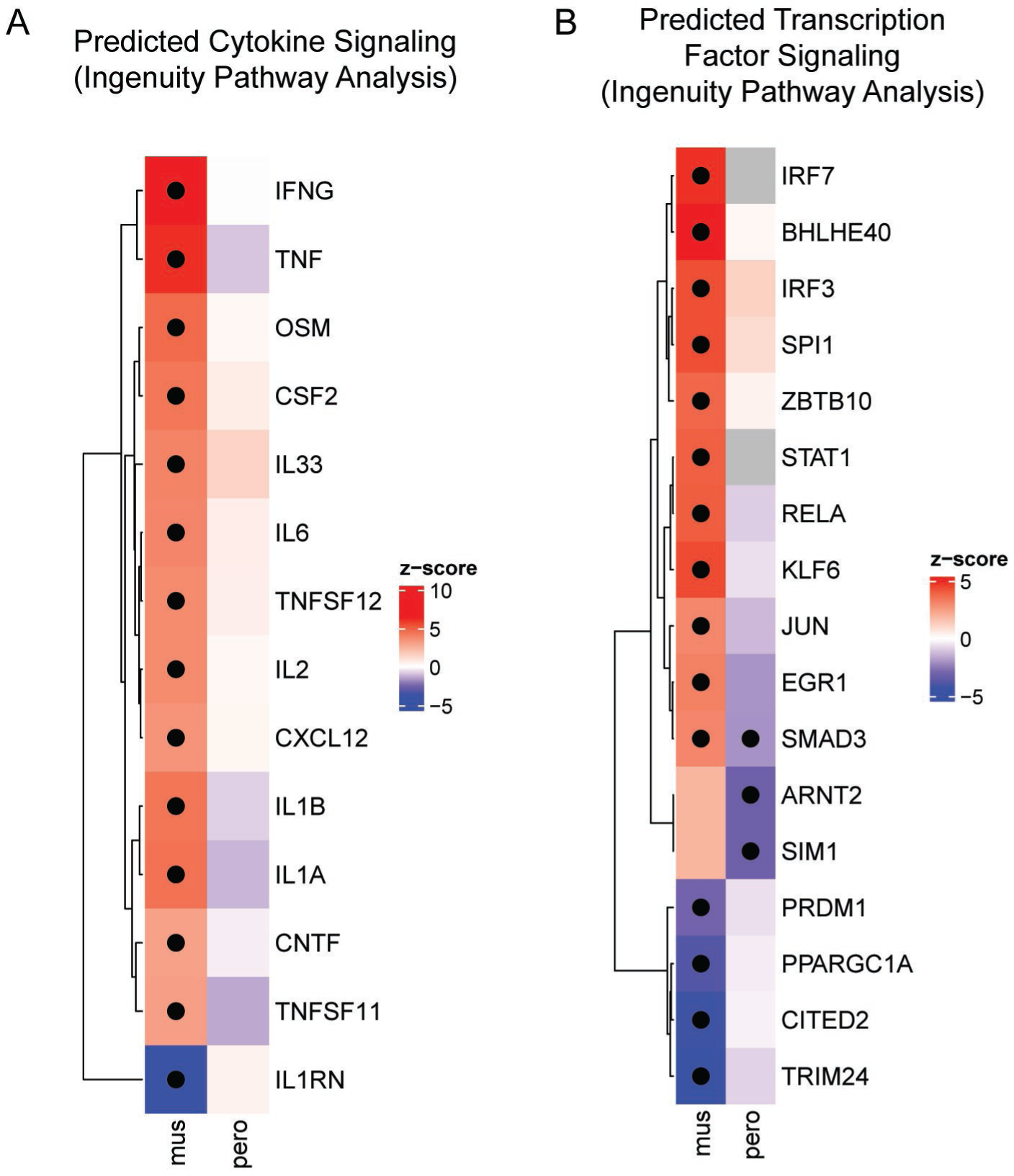
Ingenuity Pathway Analysis reveals differences in predicted pathway activation. (A) Cytokine-regulated pathways identified by QIAGEN IPA software [35] that were predicted to be differentially activated (Z score greater than or equal to 3 or less than or equal to −3) in either BALB/c mice or *P. leucopus* and had a Z score difference of 2 or greater between BALB/c and *P. leucopus*. (B) Transcription factor-regulated pathways identified by QIAGEN IPA software [35] that were predicted to be differentially activated (Z score greater than or equal to 3 or less than or equal to −3) in either BALB/c mice or *P. leucopus* and had a Z score difference of two or greater between BALB/c and *P. leucopus*. For A and B, color represents direction of effect (Red increased signaling, blue reduced signaling), dots represent a significant p-value for pathway enrichment (p<0.05).

### Ingenuity Pathway Analysis reveals differences in activity of transcription factor regulated pathways in BALB/c mice and *P. leucopus* bite sites

IPA analysis revealed 41 transcription factor regulated pathways that showed evidence of activation (Z-score ≥ 2) in BALB/c bite sites and 19 transcription factor regulated pathways with evidence of repression (Z-score ≤ −2) (**Table S2**). There were 4 transcription factor regulated pathways that showed evidence of enhanced activity in *P. leucopus* and 17 that showed evidence of repression. Of these, two transcription factor regulated pathways (ETS1 and GATA2) were predicted to be activated in both rodents.

Similar to our cytokine regulated pathway analyses, we focused our attention to transcription factor regulated pathways where we observed (1) the most significant predicted activation (Z-score > 3) or repression (Z-score < −3) in at least one of the rodent species, and (2) had at least a Z-score difference of 2 between the two species, which includes 37 pathways (**Figure 2B**). Complementing our cytokine data, we note numerous proinflammatory pathways were predicted to be activated in BALB/c but not *P. leucopus*, including IRF3 (BALB/c Z-score 4.35, *P. leucopus* Z-score 1.182) and RELA (BALB/c Z-score 4.04, *P. leucopus* Z-score −0.898). STAT1 (BALB/c Z-score 4.001) and IRF7 (BALB/c Z-score 4.851) pathways were predicted activated in BALB/c, but a Z-score could not be calculated in the *P. leucopus* dataset. Wound healing pathways, including the JUN regulated pathway, were also predicted to be activated in BALB/c but not *P. leucopus* (BALB/c Z-score 3.062, *P. leucopus* Z-score −1.292), which is notable as the AP-1 transcription factor is suppressed during nymphal tick feeding on mice [36]. Other cell proliferation and wound healing pathways also appear active specifically in BALB/c, including EGR1 (BALB/c Z-score 3.221, *P. leucopus* Z-score −1.993) and SMAD3 (BALB/c Z-score 3.072, *P. leucopus* Z-score −2.028). Transcription factor regulated pathways that were predicted suppressed in BALB/c mice but unaffected in *P. leucopus* included the HIF1A-repressor CITED2 (BALB/c Z-score −4.27, *P. leucopus* Z-score −0.282) and TRIM24 (BALB/c Z-score −4.33, *P. leucopus* Z-score −0.816).

### Comparison of larval tick bite transcriptomics with past examinations of nymphal tick bites

Recent work by Kurokawa *et al*. examined transcriptomic changes in BALB/c skin following a nymphal stage *I. scapularis* tick bite [24]. We compared our data from larval tick bites with these transcriptomic data from nymphal bite sites. Surprisingly, when we examined genes that are reported as differentially expressed in BALB/c in both studies, we see a negative correlation (r^2^=0.29, p=0.02)—with most shared differentially expressed genes that are downregulated in Kurokawa *et al.* being upregulated at the larval bite site (**Figure 3A**). Notably this includes proinflammatory factors *Il1b*, *Nlrp3*, *Ccl2*, and *Cxcl2*. In contrast, when comparing *P. leucopus* larval bite site differentially expressed genes to BALB/c nymphal bite site differentially expressed genes, a positive correlation is observed (r^2^=0.27, p<0.001): most genes are downregulated in both bite sites (**Figure 3B**). For instance, the AP-1 subunit *Fosb* was downregulated at both the nymphal BALB/c bite site and the larval *P. leucopus* bite site. Together, these results demonstrate that while we observed proinflammatory signatures in the BALB/c larval tick bite site, the BALB/c nymphal tick bite site and *P. leucopus* larval bite site appear to be more consistently anti-inflammatory.

**Figure 3:**
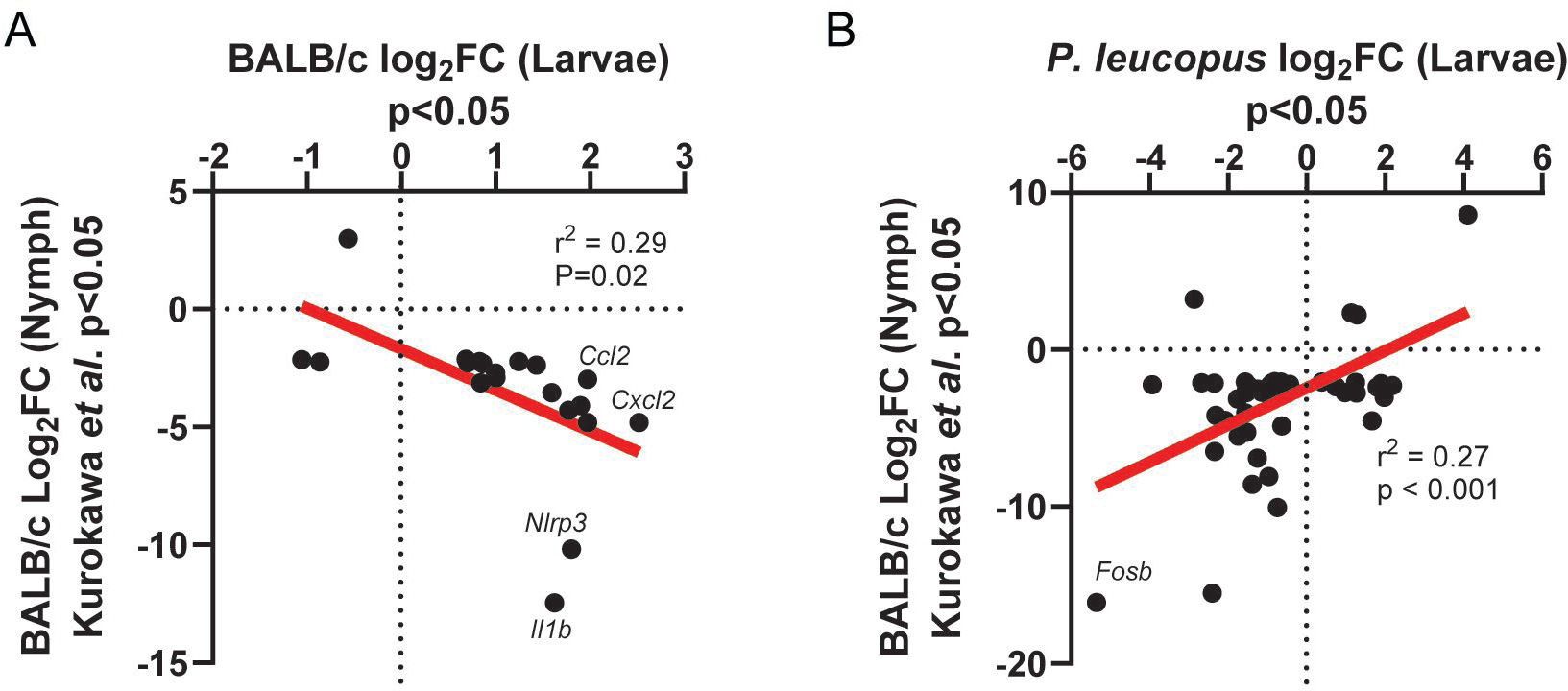
BALB/c mice, but not *P. leucopus*, display substantially different gene expression patterns following a larval tick bite compared to a BALB/c nymphal tick bite. (A,B) Comparison of differentially expressed genes (p<0.05) in the (A) BALB/c or (B) *P. leucopus* larval tick bite site compared to differentially expressed genes in the BALB/c nymphal tick bite site reported by Kurokawa et al. [24]. Statistics derived from simple linear regression and p-value describes a slope deviation from 0.

### Human larval tick bites show macroscopic and transcriptomic evidence of inflammation

Together, our data suggested that larval *I. scapularis* may be better suited at suppressing proinflammatory pathways in a native host (*P. leucopus*) than in a non-native host (*M. musculus*). This led us to hypothesize that as hosts become more evolutionarily divergent from *P. leucopus*, *I. scapularis* larvae will become less able to suppress inflammation—even on hosts where nymphs are well-described as being able to feed. To test this hypothesis, we took advantage of a human trial where patients with prior Lyme disease who had completed antibiotic therapy were exposed to 25-30 laboratory-reared larval ticks. Skin biopsies were taken before tick placement and/or after ticks were allowed to feed to repletion (NCT02446626) (**Table 1**). Unlike our rodent hosts, we routinely noted strong, macroscopic observations of inflammation at the human site of larval tick feeding (**Table 1**), though we note this may have been driven in part by the fact that these patients had previously been exposed to ticks in nature [25].

**Table 1.** Demographics of sequenced subjects.

| **Table 1. Demographics of sequenced subjects.** |  |  |  |  |  |  |  |  |
| --- | --- | --- | --- | --- | --- | --- | --- | --- |
| **Subject Code** | **Analysis Group-Feeding** | **Age** | **Sex** | **#Tick bites past 5yrs** | **Total # Fed Engorged** | **Total # Partial Fed** | **Total Ticks** | **Local reaction (definitely or probably related to the tick bite)** |
| A001* | Good feeding | 70-79 | Male | 5 | 17 | 4 | 21 | None |
| A002 | Good feeding | 70-79 | Female | 1 | 17 | 4 | 21 | Mild pruritus |
| A003 | Good feeding | 60-69 | Female | 0 | 16 | 4 | 20 | Mild pruritus |
| A004 | Good feeding | 30-39 | Male | 2 | 21 | 2 | 23 | None |
| A005 | Good feeding | 50-59 | Male | 1 | 20 | 8 | 28 | Mild pruritus |
| A006 | N/A% | 60-69 | Male | 5 | 15 | 2 | 17 | None |
| A007 | N/A% | 50-59 | Male | 1 | 12 | 3 | 15 | None |
| A009 | Good feeding | 60-69 | Female | 0 | 12 | 11 | 23 | None |
| B001* | Bad feeding | 60-69 | Female | 2 | 3 | 1 | 4 | Mild pruritus and erythema |
| B002 | Bad feeding | 70-79 | Male | 2 | 4 | 3 | 7 | Mild pruritus and moderate tenderness |
| B003* | Bad feeding | 50-59 | Male | 2 | 2 | 4 | 6 | Mild pruritus |
| B004 | Bad feeding | 20-29 | Female | 10 | 2 | 1 | 3 | Mild pruritus |
| B005** | Bad feeding | 50-59 | Male | 0 | 2 | 7 | 9 | Mild pruritus, pain and tenderness |
| B006** | N/A% | 50-59 | Male | 0 | 10 | 3 | 13 | Mild pruritus and tenderness |
| B007* | N/A% | 50-59 | Male | 3 | 7 | 0 | 7 | Vesicle at site |
| B008 | N/A% | 50-59 | Male | 1 | 4 | 10 | 14 | Mild pruritus |
| B009 | Bad feeding | 40-49 | Male | 1 | 0 | 2 | 2 | None |
| B010 | Bad feeding | 60-69 | Female | 3 | 3 | 4 | 7 | Mild pruritus |
| * These subjects had two tick placements; this information represents the first removals for B001, B004, and B007 and second removals for A001 and B003.<br>**B005 and B006 are the same subject. B006 is the second tick placement/removal.<br>%Samples with >10 but <20 recovered fed ticks were not included for good vs bad feeding analyses |  |  |  |  |  |  |  |  |

We performed RNA-sequencing on biopsies that met one of the following criteria: (1) there was a biopsy taken prior to tick placement and after tick removal (pre vs. post), (2) the post-tick removal biopsy was collected from a subject who had at least 20 total fed ticks (Good Feeding), or (3) the post-tick removal biopsy was collected from a subject who had less than 10 total fed ticks with less than half of those being fully fed ticks (Bad Feeding). There were 29 skin biopsy samples that met these criteria and were processed for RNA sequencing, from 17 participants, with 1 subject having biopsies sequenced from two xenodiagnostic procedures (**Table 1**). Human biopsies were taken at the end of tick feeding rather than at an intermediate timepoint as was used in our rodent studies.

Skin biopsies were collected from a subset of patients prior to tick placement and all patients after tick removal to examine the effects on the transcriptional profile of human skin after larval tick bites (**Figure 4A**). Principal component analysis revealed samples from bitten skin modestly clustered along the second principal component (**Figure 4B**). Differential gene expression analysis (p_adj_<0.05) revealed 4,322 differentially expressed genes (**Figure 4C, Table S3**), 2,686 of which were upregulated and 1,636 were downregulated.

**Figure 4:**
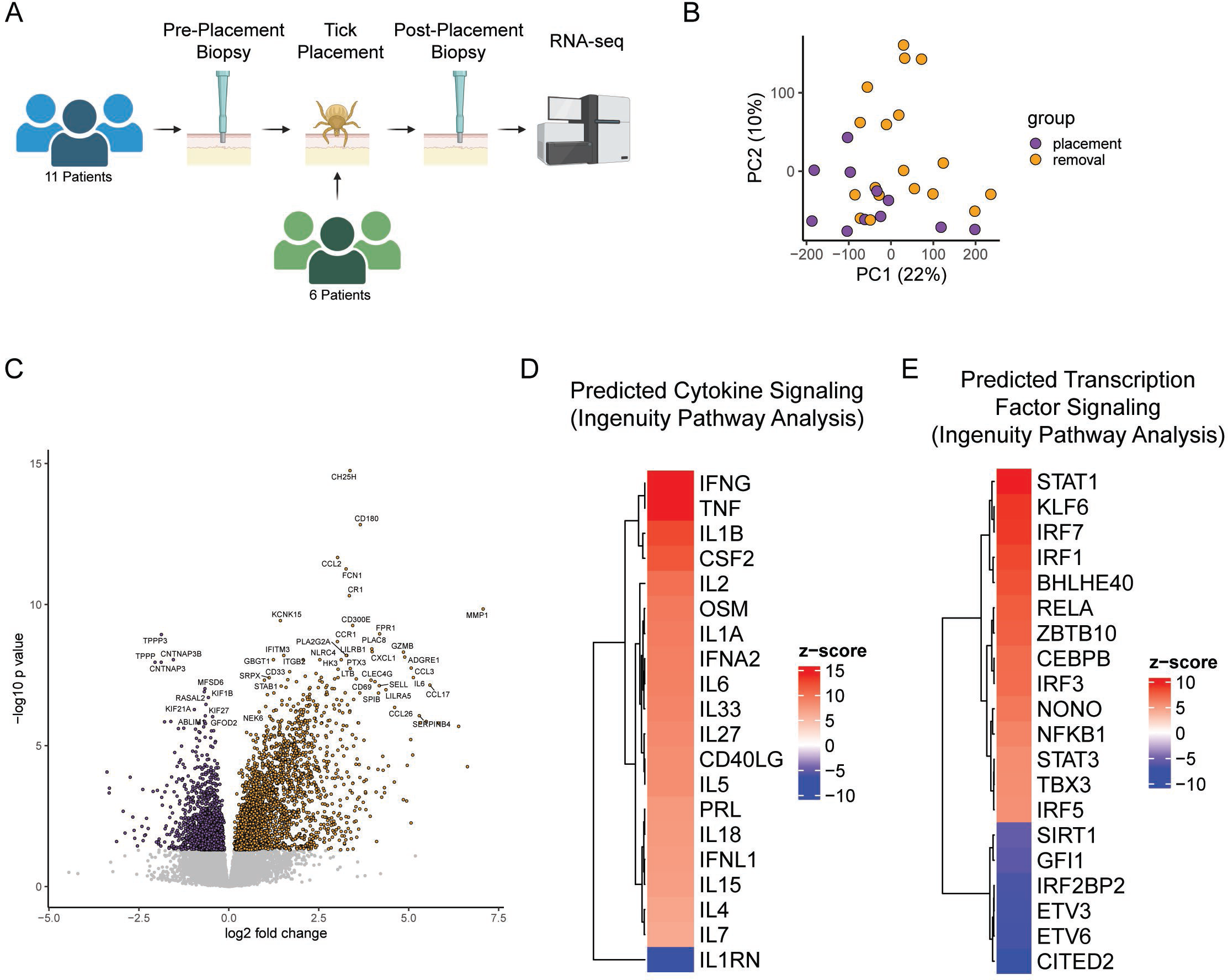
Transcriptomic analysis of the human larval *I. scapularis* bite site. (A) Schematic of patient sample collection. (B) Principal component analysis based on observed transcripts per million reads for each gene in humans. (C) Volcano plot displaying differentially expressed genes in response to the larval tick bite. Genes with p>0.05 are displayed in gray. (D,E) IPA analysis of the top 20 predicted differentially activated pathways regulated by (D) cytokines or (E) transcription factors following a larval tick bite in humans. All listed upstream regulators have a Z-score greater or equal to 2 or less than or equal to −2.

While there are many differences between our rodent and human patient studies that make direct comparison difficult, we do note that this degree of differential expression and the overall effect sizes observed are substantially larger in humans than in rodents—which aligns with our hypothesis that the response to *I. scapularis* larvae is less blunted than in natural hosts. Further, we noted robust evidence of inflammation, including significant upregulation of the macrophage chemoattractants CCL2, CCL3 and CCL4; the monocyte chemoattractant CCL8, the neutrophil chemoattractants CXCL1, CXCL2, and CXCL8, and the proinflammatory cytokines IL6, IL1B, TNF, and IL32. We noted upregulation of CD14 and CD68, markers of dendritic cells and macrophages, respectively. The adaptive immune system also appears to play a role, with B lymphocyte attractant CXCL13, T cell markers CD4 and CD69, and T cell attractants CXCL9, CXCL10, CXCL11, and CCL2 all significantly upregulated in the post-tick removal samples.

Using IPA analysis, we observed an overall pattern of elevated proinflammatory cytokine signaling in bitten human skin, including activation of TNF, IFNG, IL1, STAT3, STAT1, NfKb pathways, among others (**Table S4, Figure 4D, 4E**). We also observed suppression of IL1RN and CITED2 pathways. Notably, these pathways largely overlap with what we observed with BALB/c, suggesting that the BALB/c response to larval *I. scapularis* may be more similar to the human response than to *P. leucopus*—although the overall evidence of inflammation in the human transcriptional dataset was substantially stronger. Recent work has examined the human transcriptome in response to a nymphal *I. scapularis* tick bite [37] and observed upregulation of IL17-mediated inflammation, which we also observed in this dataset.

### Comparison of transcriptional profiles at tick bite sites between patients with high vs low percentage of tick feeding identifies a small set of differentially expressed genes

Our observations that human patients had differential abilities to support larval tick feeding led us to question if there were differences in transcriptional profiles which may be responsible for “good” (≥20 fed ticks recovered) or “bad” feeding (≤10 fed ticks recovered). Good and bad feeding did not correlate with number of reported tick exposures in the last five years (p=0.2, linear regression), pruritus (p=0.4, T test), or sex (p=0.9, T test). We compared the gene expression profiles of biopsies taken at the tick removal visit and PCA of normalized gene expression data showed no distinct separation between good and bad feeding along the first and second principal components (**Figure 5A**). Differential expression analysis revealed only 6 differentially expressed genes between feeding groups, with bad feeding as the reference group (**Figure 5B, Table S3**). There were 2 genes, DHRS2 and PM20D1, which were upregulated with a padj value cutoff of 0.05 and 4 genes, FOXCUT, MATN4, KCNJ12, and SMAD1-AS1, which were downregulated with a p_adj_ cutoff of 0.05. None of these differentially expressed genes have previously been implicated in a response to tick feeding and are not associated with an immune response.

**Figure 5:**
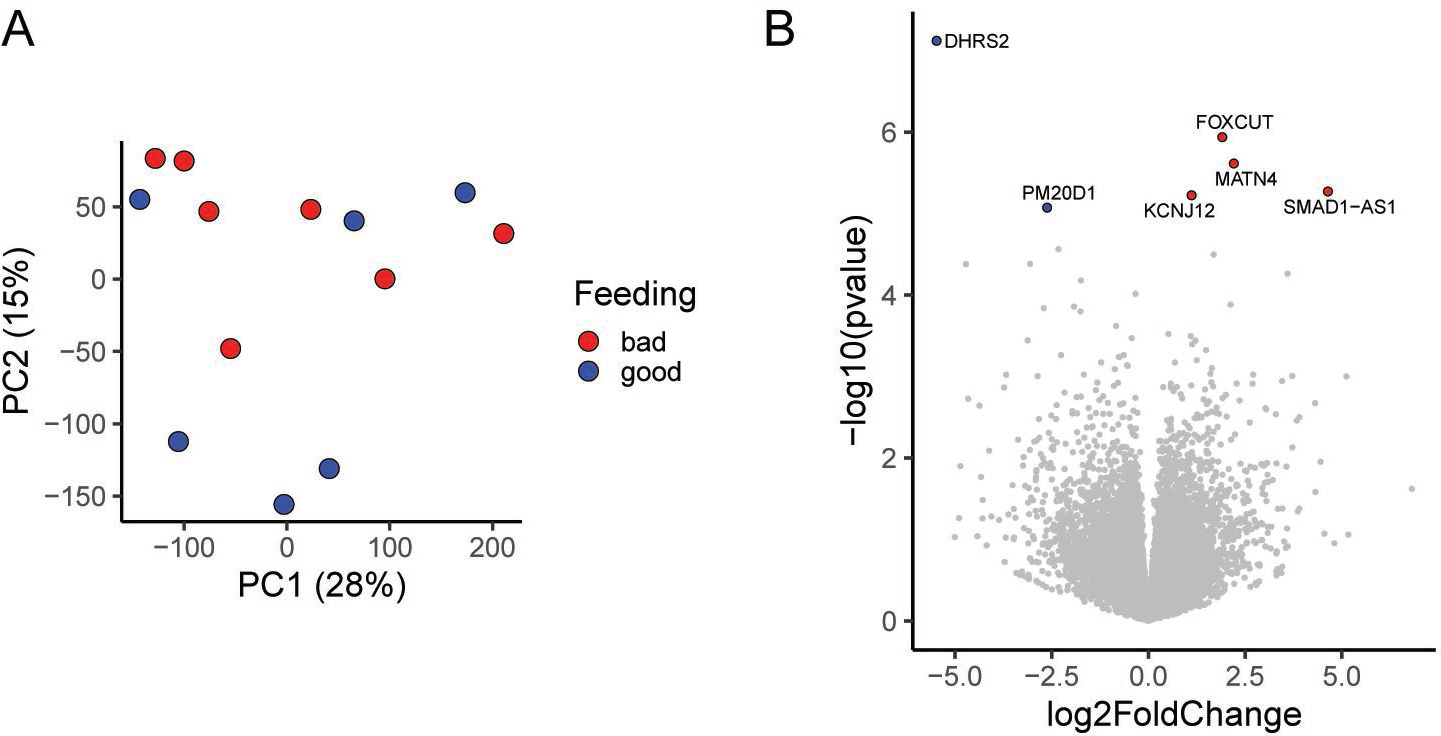
There are limited transcriptional differences between humans that had good and bad larval tick feeding. (A) Principal component analysis fails to separate individuals that had good feeding (≥20 fed ticks) or bad feeding (≤10 fed ticks) based on observed transcripts per million reads for each gene. (B) Volcano plot displaying differentially expressed genes in individuals that had good or bad feeding. Bad feeding was used as the reference dataset for differential expression. Genes with p>0.05 are displayed in gray.

## Discussion

In this study we observed divergent transcriptomic responses to bites from larval *I. scapularis* by *P. leucopus*, *M. musculus*, and humans. We specifically noted that *P. leucopus* display less evidence of skin inflammation based on the transcriptome than *M. musculus*, with humans showing far more skin inflammation than either rodent species. Importantly, however, there are several caveats to directly comparing the human results to those from the rodent studies. First, the humans participating in this study had varying levels of past tick exposure (Table 1), whereas the rodents studied were naive to tick bites, a difference which might explain the enhanced inflammatory response in human patients. Second, past work has demonstrated that the *P. leucopus* nymphal tick bite site becomes more inflamed over time [22], and thus the differences between the rodent hosts and humans may have been reduced if ticks had been allowed to feed to completion on rodent hosts, as they were on humans. However, because larvae do not leave any evidence of feeding on rodent skin, it is not possible to identify the bite site to perform the skin biopsy following completion of feeding. Despite these limitations, these data provide the first insight into comparative vertebrate immune responses to bites from larval *I. scapularis*.

We consider there to be two ways to interpret the lack of evidence of proinflammatory cytokine signatures in *P. leucopus*. The first hypothesis is that larval ticks are able to escape detection by the immune system during feeding—while *M. musculus* and humans are able to detect the larval tick bite and respond. An alternative explanation is that larval are able to exploit homeostatic immune processes and/or feedback loops (reviewed [38, 39]) in *P. leucopus* but not *M. musculus* or humans. In this second explanation the *P. leucopus* immune system could be responding to the tick bite as strongly as it does in *M. musculus* or humans but the processes that become active are very different. While we do not detect evidence of the latter in our RNA-sequencing dataset, we note that subtle changes in anti-inflammatory factors, cellular metabolism [40], or tissue resident macrophages [41] could be difficult to detect by bulk RNA-sequencing while having major impacts. This highlights the need for additional studies leveraging more sensitive tools, including single cell RNA-sequencing.

Regardless of whether larval ticks use stealth or recruitment of homeostatic processes to feed on competent hosts, there have been two non-mutually exclusive hypotheses proposed to describe host-susceptibility [6]. The first is that permissive hosts have immune systems which are intrinsically conducive/non-responsive to tick feeding. The second is that the tick is well-suited through its salivary proteins to manipulate certain host responses locally. One interpretation of our data may be that saliva at different stages of tick development (larvae vs nymph) may impact host responses differently. This could be the result of different salivary protein content or saliva abundance—particularly given the small size of larval ticks and low volume of their secreted saliva compared to nymphal ticks. The former could be impacted by differences in the tick responses to different blood sources. The composition of *I. scapularis* saliva has been found to change based on the host on which it is feeding, with *C. porcellus,* but not *M. musculus,* inducing salivary protein expression that, in turn, drives IL-4 production in murine splenocytes [42]. Nevertheless, nymphal ticks appear capable of circumventing innate immunity in *P. leucopus*, *M. musculus*, and humans, larvae appear to display some host specificity in their capacity to blunt immune responses. We do note that while we see differences in inflammation, these data should not be misinterpreted: *I. scapularis* larvae can feed on all three hosts. It will be interesting to examine whether other wildlife species display differential immune responses to larval tick feeding and whether this contributes to feeding success. Additionally, in this study BALB/c mice were used as representative *M. musculus* mice, however, additional studies may examine whether there are differences in *M. musculus* responses to larval ticks if different strains and/or outbred mice are used.

These data add to a growing number of reports [43, 44] that bring into question whether *M. musculus* is a useful model for modeling the North American *B. burgdorferi* enzootic cycle [1]—the cycle through which the spirochete transitions between vertebrate and invertebrate host in nature. While *B. burgdorferi* can cycle through *M. musculus*—particularly in Europe, the predominant reservoirs in North America are *P. leucopus* and shrews [26, 45–47]. The New World *Peromyscus* deermouse is approximately 25 million years diverged from the Eurasian *Mus* mouse [48], meaning these species have undergone substantial independent evolution and that there has been ample opportunity for *I. scapularis* to coevolve with *P. leucopus.* While previous studies have focused on differences in how these rodents interact with *B. burgdorferi* [43, 44], here we suggest that *P. leucopus* and *M. musculus* also have highly distinct cutaneous transcriptome responses to larval *I. scapularis* tick bites. Thus, while *M. musculus* has served as a very useful model for studying human Lyme disease severity and susceptibility, studies focused on the North American enzootic cycle should prioritize using natural reservoir species.

Our data comparing hosts with “good” or “bad” tick feeding do not support an association between tick feeding success and proinflammatory transcriptomic responses in humans. There are numerous alternative explanations for variable feeding success independent of host immunity. In the 4-6 days between tick placement and tick removal visits, subjects were asked to keep the dressing surrounding the ticks dry and are asked not to wear strong smelling perfumes or lotions, but subject non-compliance in any of these areas could contribute to decreases in duration of tick feeding. Additionally, larval tick handling by physicians during the clinical trial was variable, and accidental damage of ticks during placement on patients is possible. Further, our protocol for collecting the skin biopsies did not capture the participants with the worst feeding, as skin biopsies were only collected from subjects with at least one fed tick.

In conclusion, these findings provide new insights into divergent host responses to larval *I. scapularis* tick feeding. We anticipate that these findings may serve as a launching point to identify potential pharmacologic interventions to disrupt larval tick feeding in nature. Disrupting this stage of the *Ixodes* life cycle not only prevents development of ticks and further reproduction, but blocks tick acquisition of enzootic-cycling pathogens and eventual spillover into humans. Further studies may examine how this skin inflammation affects the tick-host-pathogen interface during enzootic cycling. Curiously, despite observing a stronger proinflammatory response here at the larval tick-*M. musculus* interface, past work demonstrates that *B. burgdorferi* is actually more readily transmitted from *M. musculus* to larval *I. scapularis* than from *P. leucopus* to *I. scapularis* [43, 49]. Thus, the enhanced inflammation we observed at the *M. musculus* larval tick bite apparently does not impair transmission of the spirochete to new invertebrate hosts.

## Acknowledgments

This study was funded by the National Institutes of Health (JSB was supported by F32AI179104; LTH by R01AI152210, U01AI109656, and R21AI146841; SRT by R01AI137424 and U01AI109656). It was also supported in part by the Division of Intramural Research, National Institute of Allergy and Infectious Diseases, National Institutes of Health (SPT, AM). We thank all members of the Hu lab and Joao Pedra for helpful discussions throughout the preparation of this manuscript. We thank Tufts Comparative Medical Services for enabling these experiments. RNA sequencing analyses were made possible by the Tufts High Performance Computing Cluster (https://it.tufts.edu/high-performance-computing). Figures 1A and 4A were generated using BioRender.com.

## Supplemental Materials Legends

Table S1: DESeq2 analysis of BALB/c *M. musculus* and *P. leucopus* gene expression following larval tick bite.

Table S2: QIAGEN IPA analysis of BALB/c *M. musculus* and *P. leucopus* signaling following larval tick bite.

Table S3: DESeq2 analysis of human gene expression following larval tick bite. Data contain two analyses: Pre- and Post-tick placement and good (>20 fed ticks) vs bad (<10 fed ticks) feeding.

Table S4: QIAGEN IPA analysis of human signaling following larval tick bite.

## Materials and Methods

### Rodent Maintenance, Use, and Ethical Statement

All animal procedures were approved by the Tufts University-Tufts Medical Center Institutional Animal Care and Use Committee (IACUC, Protocol #B2021-84). Euthanasia was performed in accordance with guidelines provided by the American Veterinary Medical Association (AVMA) and was approved by the Tufts University IACUC. Rodents were maintained by the Tufts Comparative Medicine Services. The *P. leucopus* colony was started by Dr. Sam Telford using wild captured rodents from the northeastern and midwestern United States. The colony has been closed since 1994, held in microisolator cages, and is specific pathogen free (regular sentinel testing). BALB/c mice were obtained from Charles River Laboratories (Strain Code 028).

Equal numbers of male and female rodents were used at the beginning of the experiment, however, variable rates of successful recovery of ticks and/or RNA led to sex-imbalance: Two male and three female *P. leucopus* were used in the study; One male and three female BALB/c mice were used.

### Rodent Tick Infestation

A heated 1:4 mixture of melted beeswax to rosin gum mixture was used to attach a modified microcentrifuge tube lined with mesh between the shaved shoulder blades of each rodent. Ten *I. scapularis* larvae (National Tick Research and Education Resource, Oklahoma State University) were placed inside the mesh cap. Mice were singly housed in cages surrounded by a water moat for 48 hours. The tick containment chamber was carefully removed, and a 2-mm punch biopsy was taken surrounding a single feeding tick and transferred immediately to RNAlater, stored overnight at room temperature, and then frozen at –80°C until use. Biopsies were intentionally restricted to regions away from the edge of the tick containment chamber where damage from removal could skew results. A second biopsy was taken on the outside of the tick containment chamber as a matched control.

### Human subjects

The subjects described in this study were enrolled in the Xenodiagnosis After Antibiotic Treatment for Lyme Disease clinical study (NCT02446626). The study was approved by the Institutional Review Boards at each center and written informed consent was obtained from all participants. Participants were enrolled at Tufts University in Boston, Massachusetts; National Institutes of Health in Bethesda, Maryland; Mansfield Family Practice in Storrs, Connecticut, and Stony Brook University in Stony Brook, New York. Sequencing of banked biopsy samples was approved by Tufts University Institutional Review Board.

### Human Tick preparation, placement, and removal

Larval *Ixodes scapularis* ticks were reared at a central facility and provided to each of the testing centers. Tick placement and removal was performed as described by Turk et al [50]. For some participants in the Post Treatment Lyme Disease Syndrome group, punch skin biopsies were taken at a control site prior to tick placement and for all participants, biopsies were taken at the completion of tick feeding, at the site of a tick bite.

### Rodent RNA Library Preparation and RNA sequencing

After all rodent samples were collected as described above, samples were thawed on ice, transferred to QIAzol, and bead beaten with a 6.35 mm chrome steel bead (Biospec Products) for two cycles of two minutes each at an oscillation frequency of 30/second using a TissueLyser II (Qiagen). RNA was extracted using the miRNeasy mini kit (Qiagen), with the modification that two RWT washes were performed. gDNA was digested using TURBO DNase (Invitrogen) and RNA was repurified using the RNeasy MinElute Cleanup Kit (Qiagen). Purified RNA was submitted to Azenta Life Sciences/Genewiz for library preparation and sequencing (Illumina HiSeq 2×150bp).

### Human RNA Library Preparation and RNA sequencing

2mm skin biopsies from subjects were stored in RNAlater overnight at room temperature and then frozen at –80°C until use. Samples were homogenized under liquid nitrogen using a mortar and pestle. Tissues were then resuspended in 1 mL of lysis buffer from the RNeasy Fibrous Tissue Mini Kit (Qiagen) and processed as per the manufacturer’s protocol. Ribosomal RNA was depleted and RNA-seq libraries were prepared by Tufts University Core Facility. Sequencing of 50bp single end reads was performed on a HiSeq 2500.

### RNA-seq Analysis

Reads were mapped to *P. leucopus* (GCF_004664715.2) [44], *M. musculus* (GRCm39), or human genomes (GRCh38.98) using STAR version 2.6.1 [51] and gene expression was summarized using RSEM version 1.3.1 [52]. Principal component analysis was performed using the R Bioconductor PCAtools package (R version 4.3.1) [53]. Differential expression analysis was performed using DEseq2 (R version 4.3.1) [54]. Genes with low expression were removed based on their normalized read counts. Genes with greater than 10 read counts were kept. DEGs were identified using an adjusted p-value cutoff of 0.05. Genes with no expression and T cell receptor genes were removed prior to plotting. Heatmaps and clustering, based on Euclidean distance, were generated using the ComplexHeatmap package (R version 4.3.1) [55]. Pathway analysis was performed with the use of QIAGEN Ingenuity Pathway Analysis (QIAGEN Inc., https://digitalinsights.qiagen.com/IPA) [35]. IPA used differentially expressed genes from the DESeq2 differential expression analysis with p_adj_≤0.05.

### Data Availability

Rodent sequencing data are deposited in GEO (GSE266088) and data from the human clinical trial will be available in dbGaP (phs003314.v1.p1).

## References

1. Radolf, J.D., et al., Of ticks, mice and men: understanding the dual-host lifestyle of Lyme disease spirochaetes. Nat Rev Microbiol, 2012. 10(2): p. 87–99.

2. Kugeler, K.J., et al., Estimating the Frequency of Lyme Disease Diagnoses, United States, 2010-2018. Emerg Infect Dis, 2021. 27(2): p. 616–619.

3. Barbour, A.G., Infection resistance and tolerance in Peromyscus spp., natural reservoirs of microbes that are virulent for humans. Semin Cell Dev Biol, 2017. 61: p. 115–122.

4. Eisen, R.J. and L. Eisen, The Blacklegged Tick, Ixodes scapularis: An Increasing Public Health Concern. Trends Parasitol, 2018. 34(4): p. 295–309.

5. Simo, L., et al., The Essential Role of Tick Salivary Glands and Saliva in Tick Feeding and Pathogen Transmission. Front Cell Infect Microbiol, 2017. 7: p. 281.

6. Ribeiro, J.M., Role of saliva in tick/host interactions. Exp Appl Acarol, 1989. 7(1): p. 15–20.

7. Wikel, S., Ticks and tick-borne pathogens at the cutaneous interface: host defenses, tick countermeasures, and a suitable environment for pathogen establishment. Front Microbiol, 2013. 4: p. 337.

8. Kazimirova, M. and I. Stibraniova, Tick salivary compounds: their role in modulation of host defences and pathogen transmission. Front Cell Infect Microbiol, 2013. 3: p. 43.

9. Francischetti, I.M., et al., The role of saliva in tick feeding. Front Biosci (Landmark Ed), 2009. 14: p. 2051–88.

10. Piesman, J., J.R. Oliver, and R.J. Sinsky, Growth kinetics of the Lyme disease spirochete (Borrelia burgdorferi) in vector ticks (Ixodes dammini). Am J Trop Med Hyg, 1990. 42(4): p. 352–7.

11. De Silva, A.M. and E. Fikrig, Growth and migration of Borrelia burgdorferi in Ixodes ticks during blood feeding. Am J Trop Med Hyg, 1995. 53(4): p. 397–404.

12. Ribeiro, J.M., et al., Dissemination and salivary delivery of Lyme disease spirochetes in vector ticks (Acari: Ixodidae). J Med Entomol, 1987. 24(2): p. 201–5.

13. Kitsou, C., E. Fikrig, and U. Pal, Tick host immunity: vector immunomodulation and acquired tick resistance. Trends Immunol, 2021. 42(7): p. 554–574.

14. Ramamoorthi, N., et al., The Lyme disease agent exploits a tick protein to infect the mammalian host. Nature, 2005. 436(7050): p. 573–7.

15. Schuijt, T.J., et al., The tick salivary protein Salp15 inhibits the killing of serum-sensitive Borrelia burgdorferi sensu lato isolates. Infect Immun, 2008. 76(7): p. 2888–94.

16. Anguita, J., et al., Salp15, an ixodes scapularis salivary protein, inhibits CD4(+) T cell activation. Immunity, 2002. 16(6): p. 849–59.

17. Schuijt, T.J., et al., A tick mannose-binding lectin inhibitor interferes with the vertebrate complement cascade to enhance transmission of the lyme disease agent. Cell Host Microbe, 2011. 10(2): p. 136–46.

18. Nguyen, T.T., et al., A tick saliva serpin, IxsS17 inhibits host innate immune system proteases and enhances host colonization by Lyme disease agent. PLoS Pathog, 2024. 20(2): p. e1012032.

19. Tang, X., et al., A tick C1q protein alters infectivity of the Lyme disease agent by modulating interferon gamma. Cell Rep, 2022. 41(8): p. 111673.

20. Kurokawa, C., et al., Interactions between Borrelia burgdorferi and ticks. Nat Rev Microbiol, 2020. 18(10): p. 587–600.

21. Keirans, J.E., et al., Ixodes (Ixodes) scapularis (Acari:Ixodidae): redescription of all active stages, distribution, hosts, geographical variation, and medical and veterinary importance. J Med Entomol, 1996. 33(3): p. 297–318.

22. Anderson, J.M., et al., Ticks, Ixodes scapularis, Feed Repeatedly on White-Footed Mice despite Strong Inflammatory Response: An Expanding Paradigm for Understanding Tick-Host Interactions. Front Immunol, 2017. 8: p. 1784.

23. Krause, P.J., et al., Dermatologic changes induced by repeated Ixodes scapularis bites and implications for prevention of tick-borne infection. Vector Borne Zoonotic Dis, 2009. 9(6): p. 603–10.

24. Kurokawa, C., et al., Repeat tick exposure elicits distinct immune responses in guinea pigs and mice. Ticks Tick Borne Dis, 2020. 11(6): p. 101529.

25. Burke, G., et al., Hypersensitivity to ticks and Lyme disease risk. Emerg Infect Dis, 2005. 11(1): p. 36–41.

26. Goethert, H.K., et al., Host-utilization differences between larval and nymphal deer ticks in northeastern U.S. sites enzootic for Borrelia burgdorferi sensu stricto. Ticks Tick Borne Dis, 2023. 14(6): p. 102230.

27. Piesman, J. and A. Spielman, Host-Associations and Seasonal Abundance of Immature Ixodes dammini1 in Southeastern Massachusetts2. Annals of the Entomological Society of America, 1979. 72(6): p. 829–832.

28. Godsey, M.S., Jr., et al., Lyme disease ecology in Wisconsin: distribution and host preferences of Ixodes dammini, and prevalence of antibody to Borrelia burgdorferi in small mammals. Am J Trop Med Hyg, 1987. 37(1): p. 180–7.

29. Kollars, T.M., Jr., et al., Seasonal activity and host associations of Ixodes scapularis (Acari: Ixodidae) in southeastern Missouri. J Med Entomol, 1999. 36(6): p. 720–6.

30. Ginsberg, H.S., et al., Selective Host Attachment by Ixodes scapularis (Acari: Ixodidae): Tick-Lizard Associations in the Southeastern United States. J Med Entomol, 2022. 59(1): p. 267–272.

31. Davidar, P., M. Wilson, and J.M. Ribeiro, Differential distribution of immature Ixodes dammini (Acari: Ixodidae) on rodent hosts. J Parasitol, 1989. 75(6): p. 898–904.

32. Rollend, L., D. Fish, and J.E. Childs, Transovarial transmission of Borrelia spirochetes by Ixodes scapularis: a summary of the literature and recent observations. Ticks Tick Borne Dis, 2013. 4(1-2): p. 46–51.

33. Xu, G., et al., Human-Biting Ixodes Ticks and Pathogen Prevalence from California, Oregon, and Washington. Vector Borne Zoonotic Dis, 2019. 19(2): p. 106–114.

34. Marques, A., et al., Xenodiagnosis to detect Borrelia burgdorferi infection: a first-in-human study. Clin Infect Dis, 2014. 58(7): p. 937–45.

35. Kramer, A., et al., Causal analysis approaches in Ingenuity Pathway Analysis. Bioinformatics, 2014. 30(4): p. 523–30.

36. Marnin, L., et al., Tick extracellular vesicles impair epidermal homeostasis through immune-epithelial networks during hematophagy. bioRxiv, 2023.

37. Tang, X., et al., Bulk and single-nucleus RNA sequencing highlight immune pathways induced in individuals during an Ixodes scapularis tick bite. Infect Immun, 2023. 91(11): p. e0028223.

38. Medzhitov, R., The spectrum of inflammatory responses. Science, 2021. 374(6571): p. 1070–1075.

39. Meizlish, M.L., et al., Tissue Homeostasis and Inflammation. Annu Rev Immunol, 2021. 39: p. 557–581.

40. Eming, S.A., P.J. Murray, and E.J. Pearce, Metabolic orchestration of the wound healing response. Cell Metab, 2021. 33(9): p. 1726–1743.

41. Zhao, J., I. Andreev, and H.M. Silva, Resident tissue macrophages: Key coordinators of tissue homeostasis beyond immunity. Sci Immunol, 2024. 9(94): p. eadd1967.

42. Narasimhan, S., et al., Host-specific expression of Ixodes scapularis salivary genes. Ticks Tick Borne Dis, 2019. 10(2): p. 386–397.

43. Bourgeois, J.S., et al., Comparative reservoir competence of Peromyscus leucopus and mouse models for Borrelia burgdorferi B31. bioRxiv, 2023: p. 2023.09.28.559638.

44. Long, A.D., et al., The genome of Peromyscus leucopus, natural host for Lyme disease and other emerging infections. Sci Adv, 2019. 5(7): p. eaaw6441.

45. LoGiudice, K., et al., The ecology of infectious disease: effects of host diversity and community composition on Lyme disease risk. Proc Natl Acad Sci U S A, 2003. 100(2): p. 567–71.

46. Goethert, H.K., et al., Retrotransposon-Based Blood Meal Analysis of Nymphal Deer Ticks Demonstrates Spatiotemporal Diversity of Borrelia burgdorferi and Babesia microti Reservoirs. Appl Environ Microbiol, 2021. 87(2).

47. Tsao, J.I., et al., An ecological approach to preventing human infection: vaccinating wild mouse reservoirs intervenes in the Lyme disease cycle. Proc Natl Acad Sci U S A, 2004. 101(52): p. 18159–64.

48. Steppan, S., R. Adkins, and J. Anderson, Phylogeny and divergence-date estimates of rapid radiations in muroid rodents based on multiple nuclear genes. Syst Biol, 2004. 53(4): p. 533–53.

49. Hanincova, K., et al., Fitness variation of Borrelia burgdorferi sensu stricto strains in mice. Appl Environ Microbiol, 2008. 74(1): p. 153–7.

50. Turk, S.P., A. Eschman, and A. Marques, A Model for Experimental Exposure of Humans to Larval Ixodes scapularis Ticks. J Vis Exp, 2023(202).

51. Dobin, A., et al., STAR: ultrafast universal RNA-seq aligner. Bioinformatics, 2013. 29(1): p. 15–21.

52. Li, B. and C.N. Dewey, RSEM: accurate transcript quantification from RNA-Seq data with or without a reference genome. BMC Bioinformatics, 2011. 12: p. 323.

53. Blighe K, L.A., PCAtools: Everything Principal Components Analysis. 2023.

54. Love, M.I., W. Huber, and S. Anders, Moderated estimation of fold change and dispersion for RNA-seq data with DESeq2. Genome Biol, 2014. 15(12): p. 550.

55. Gu, Z., R. Eils, and M. Schlesner, Complex heatmaps reveal patterns and correlations in multidimensional genomic data. Bioinformatics, 2016. 32(18): p. 2847–9.

